# Resolving Immune Lineage and Cell-State Heterogeneity in Human PBMCs via Mass Spectrometry-Based Single-Cell Proteomics

**DOI:** 10.64898/2026.07.29.741525

**Authors:** Samantha A. O’Connor, Romell B. Gletten, Ritin Sharma, Zoe N. Jensen, Brooke Lovell, Krystine Garcia-Mansfield, Lucy Y. Ghoda, Bin Zhang, David E. Frankhouser, Russell C. Rockne, Jeffrey M. Trent, Guido Marcucci, Patrick Pirrotte

## Abstract

Single-cell proteomics (SCP) currently lacks validated benchmarking standards, and cell annotation often relies on transcriptomic proxies. Unsupervised clustering offers a proxy-free alternative, but its success depends on biological signal outweighing technical variation. In homogeneous samples this is achievable, but in heterogeneous populations, where closely related cell types differ only subtly, technical variation can dominate the clustering and obscure the biology needed for annotation. To address this, we developed an integrated experimental and computational pipeline for protein-level cell annotation and applied it to human PBMCs as an immune-cell test case. We isolated T cells, B cells, monocytes, and NK cells by negative-selection sorting to build a high-fidelity reference. In parallel, unsorted PBMCs from the same donor were processed on a cellenONE and acquired using label-free DIA on an Orbitrap Astral Zoom. Using the labeled reference dataset, we systematically benchmarked normalization, imputation, and clustering methods to assess their effect on cell-type separation. Unsupervised analysis resolved functional subpopulations within each lineage, and a probabilistic SCP classifier trained on these annotations identified the corresponding cell types and states in the unsorted PBMC fraction, validating the pipeline on unenriched, heterogeneous samples. Together, this work delivers an analytically benchmarked SCP workflow that resolves immune lineage and cell-state heterogeneity in human PBMCs and provides a classifier-ready, protein-level reference for immune-cell assignment.

## INTRODUCTION

Peripheral blood mononuclear cells (PBMCs) are circulating myeloid and lymphoid cells that mediate immune surveillance and coordinate innate and adaptive immunity (Parkin and Cohen 2001; Marshall et al. 2018). In healthy individuals, PBMCs primarily comprise T cells (50-77%), monocytes (10-20%), natural killer (NK) cells (5-20%), B cells (5-10%), and dendritic cells (1-2%) (Kleiveland 2015). Each major cell type can be further divided into functionally specialized subpopulations with distinct roles in host defense and immune homeostasis (Villani et al. 2017; Papalexi and Satija 2018; Fang et al. 2018; Hao et al. 2021).

Flow cytometry, cytometry by time of flight (CyTOF), and single-cell RNA sequencing (scRNA-seq) represent invaluable tools for molecular classification of PBMCs enabling the identification of major immune cell types and refined cell subpopulations (Zheng et al. 2017; Higdon et al. 2024) while providing insight into their specialized immune functions (Villani et al. 2017; Papalexi and Satija 2018; Hao et al. 2021). Amongst these technologies, scRNA-seq has emerged as the current standard for PBMC cell type and subpopulation identification as a function of comprehensive transcriptomics coverage to resolve cellular identity and transcriptional state (Butler et al. 2018; Stuart et al. 2019; Hao et al. 2021). Despite these advances, proteins remain the primary functional mediators of cellular processes (Vogel and Marcotte 2012; Kelly 2020) and the principal molecular targets of therapeutic intervention (Santos et al. 2017; Ghadermarzi et al. 2019). Furthermore, protein abundance generally provides a more direct measure of cellular phenotype (Vogel and Marcotte 2012; Wu et al. 2026) as mRNA and protein expression exhibit only moderate concordance because of post-transcriptional, translational, and post-translational regulation (Liu et al. 2016; Leduc et al. 2025; Wu et al. 2026). Consequently, direct measurement of the PBMC proteome at single-cell resolution would provide complementary functional information that cannot be fully inferred from transcriptomic analyses alone.

Though nascent in its application, single-cell proteomics (SCP) is a viable strategy for cell subpopulation resolution and cell state identification (Leduc et al. 2024; Fulcher et al. 2024; Bubis et al. 2025; Leduc et al. 2025). SCP-based profiling of immune cell types and their corresponding proteomes represents a promising approach for biomarker discovery as PBMCs are an easily accessible, minimally invasive biospecimen routinely collected in clinical settings. Furthermore, alterations in immune cell composition and activity frequently accompany the earliest stages of infection (Medzhitov 2007), inflammation (Medzhitov 2008; L. Chen et al. 2018), cancer (Fridman et al. 2017; Binnewies et al. 2018; Hanahan 2022), and autoimmune disease (Davidson and Diamond 2001; Rosenblum et al. 2015). Thus, we sought to develop an experimental workflow and computational pipeline to resolve PBMC cell types and subpopulations via SCP. We isolated T cells, B cells, monocytes and NK cells from healthy human PBMCs by negative-selection magnetic activated cell sorting (MACS), dispensed single cells for on-plate processing using the cellenONE platform, and analyzed them by label-free data-independent (DIA) acquisition liquid chromatography tandem mass spectrometry (LC-MS/MS) on an Orbitrap Astral Zoom at 70 samples per day yielding a reference set of sorted immune-cell proteomes. In parallel, we acquired the proteomes of 887 mixed, unsorted PBMCs from the same donor material, and used the sorted, annotated proteomes to classify PBMC cell types and subpopulations in the unsorted fraction at single-cell resolution. Lastly, we performed scRNA-seq on the unsorted fraction to assess single-cell mRNA-protein concordance and to validate our SCP-based cell-lineage classifications. This study establishes a workflow for high-depth SCP characterization of PBMCs and provides a foundation for future clinical applications in which PBMCs, and leukocytes more broadly, can be profiled to identify disease-associated proteomics signatures, monitor therapeutic response, and facilitate biomarker discovery.

## RESULTS

### High-throughput SCP workflow resolves major PBMC lineages

We established a reference baseline for single-cell immune proteomes by isolating four major peripheral blood mononuclear cell (PBMC) lineages: T cells, B cells, natural killer (NK) cells, and monocytes, using negative magnetic selection (**Figure 1A**). Negative selection avoids the antibody-induced signaling and surface receptor engagement that can perturb the native cellular proteome (Hornschuh et al. 2022; Brunner et al. 2022; Ctortecka et al. 2024). To evaluate the workflow on both discrete populations and mixed matrices, we isolated and dispensed single cells from the sorted lineages and from unsorted, mixed PBMCs alongside cell-free negative controls into 384-well plates using the cellenONE platform for on-plate digestion (**Figure 1B**). Samples were acquired on an Orbitrap Astral Zoom mass spectrometer at a throughput of 70 samples per day (SPD), yielding a comprehensive discovery dataset of 2,427 (including 1,540 sorted; 887 unsorted) single-cell proteomes. Although absolute cell diameters were uniformly offset by the cellenONE in-flight optical system (Leduc et al. 2024), the expected proportional scaling and lineage-distinct diameter distributions confirmed cell integrity in both the sorted and unsorted populations (**Figure 2A-B**). We first focused on the 1,540 sorted cells, searching them together as a single cohort to establish the reference baseline. Within this cohort we identified a total of 3,670 protein groups with a median of 1,049 protein groups per cell, compared with a median of 14 in the cell-free negative controls. This ∼78-fold contrast demonstrates the sensitivity of the workflow and provided the foundation for the downstream quality-control (QC) thresholds described below.

**Figure 1.**
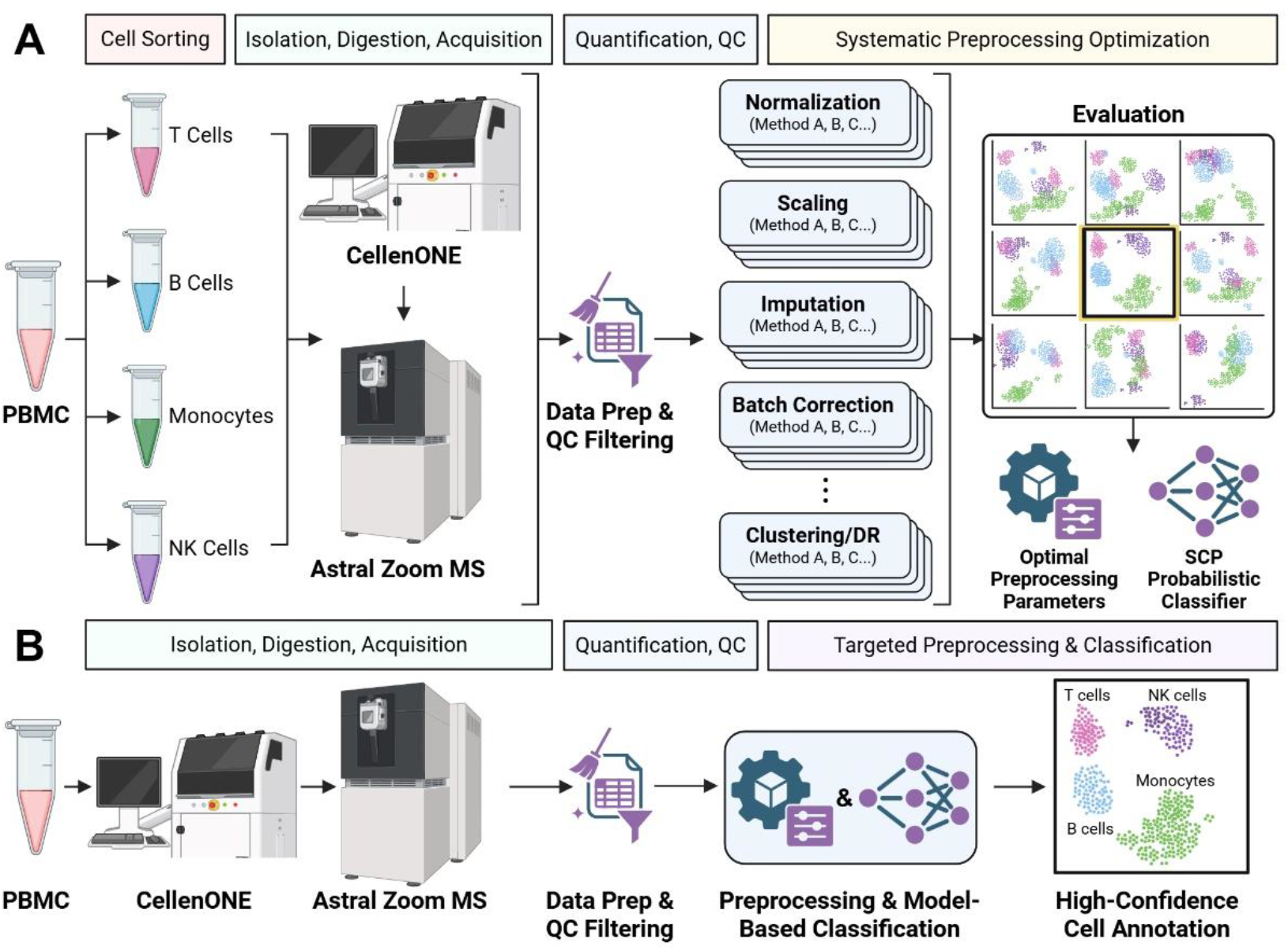
Experimental and computational workflow for single-cell proteomics of human PBMCs. **A**. Sorted PBMC lineages were processed by SCP and used to benchmark preprocessing and train a probabilistic classifier. **B**. Unsorted PBMCs from the same donor were processed using the optimized pipeline and classified for high-confidence cell annotation. Created with BioRender.com

**Figure 2.**
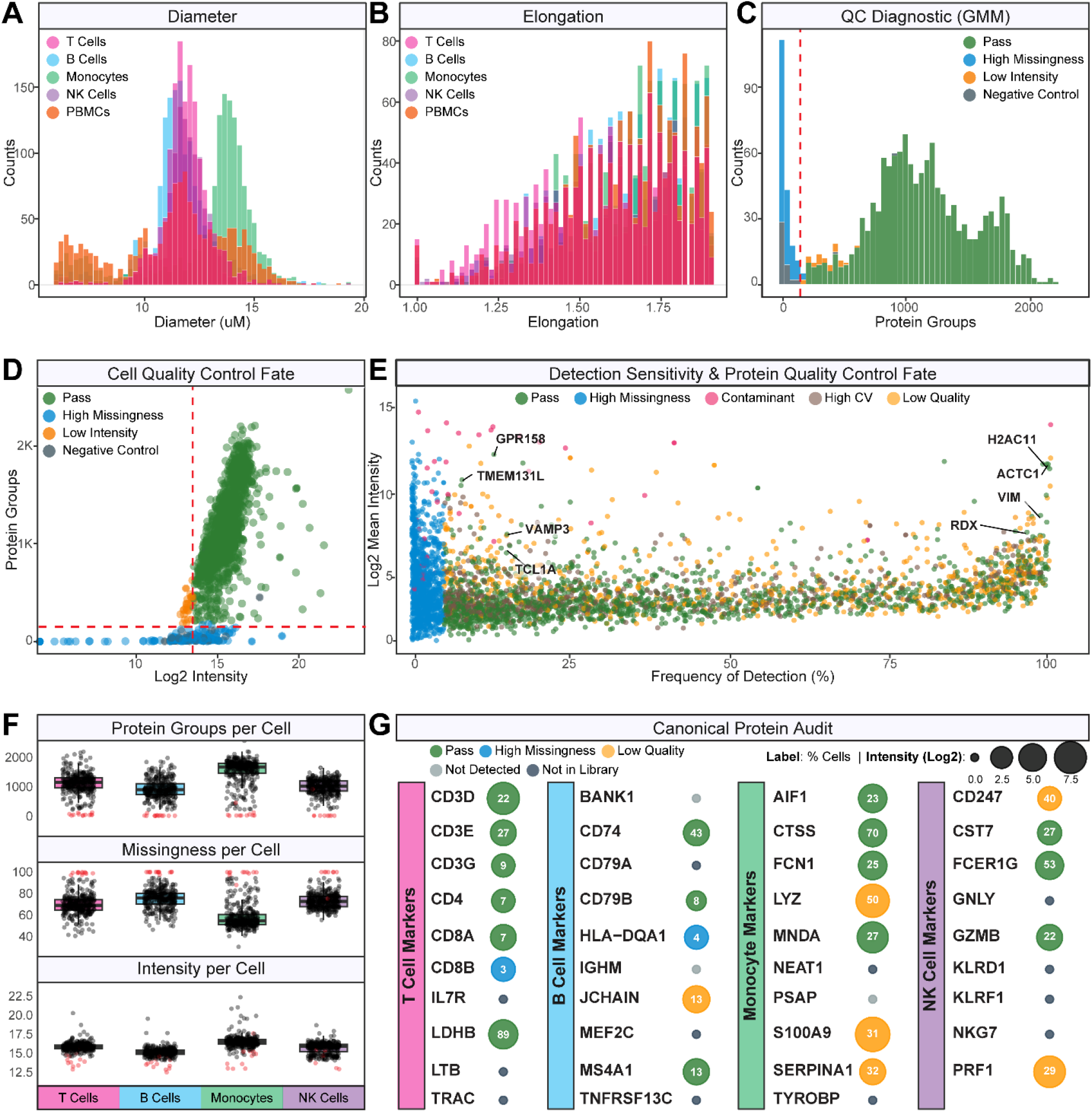
Quality control of single-cell proteomes. **A-B**. Cell diameter and elongation (obtained from the cellenONE) distributions across sorted lineages and unsorted PBMCs. **C**. Gaussian mixture model (G=3) distinguished true single cells from background by protein group count and (**D**) intensity vs. protein groups. Cells are colorized by QC fate. **E**. Log2 intensity versus detection frequency. Proteins colorized by QC fate. **F**. Protein groups, missingness, and intensity per cell across the four immune lineages. **G**. Canonical marker audit showing detection status, percentage of cells expressing, and mean log2 intensity.

### Data-driven QC defines true single-cell proteomes and marker detectability

To ensure data quality without relying on arbitrary filter cutoffs, we implemented a data-driven QC pipeline operating at both the cell and protein levels. We first fit a Gaussian mixture model (GMM), anchored to our cell-free negative controls (n = 44), to statistically define the ambient background across the sorted cohort (**Figure 2C-D**). This framework established guardrails to distinguish true single-cell signal from background noise. At the cell level, the pipeline evaluated integrity using cumulative-intensity floors, missingness, and background-probability distribution (**Figure 2D**). Of the 1,540 sorted cells, the GMM flagged 148 for high missingness (missing values for ≥ 95% of proteins) and 26 for falling below the intensity floor (**Figure S1A**). Flagged cells were excluded from normalization and downstream analysis unless manually rescued, leaving 1,322 single cells that passed QC. Retained cells aligned closely with expected biology: monocytes, being substantially larger than lymphoid lineages, showed a higher protein density (median of 1,671 protein groups per cell) and a corresponding shift toward higher MS intensities than T, B, and NK cells (median of 1,014 protein groups per cell; **Figure 2F**, see also **Figure S1C**).

Parallel QC at the protein level isolated technical artifacts from true biological features. After strict removal of common contaminants, quantitative thresholds flagged 2,166 proteins collectively for extreme missingness (>95%), high variability, and low signal-to-noise ratio, and 1,382 proteins were categorized as “not detected in any cell” (**Figure S1B**). This last group represents a known artifact of multi-stage library transfer in single-cell DIA workflows. While these protein groups were confidently detected during batch-level directDIA library generation, their underlying low-abundance precursors fell below the limit of detection in individual single-cell spectra, failing the retention-time alignment and q-value thresholds during the global library search (**Figure 2E**). Despite this protein-level filtering, the lineage-associated depth difference was preserved: monocytes retained a higher median protein count than the lymphoid lineages (896 vs. 518 protein groups per cell; **Figure S1D**), confirming that size-associated depth effect is not an artifact of the protein filter. By prospectively auditing canonical PBMC markers within these categories, we traced where and why key markers dropped out, allowing us to pre-emptively assess their viability for downstream analysis and cell annotation. A subset of canonical markers, including the T-cell marker IL7R, the B-cell marker CD79A, the monocyte marker NEAT1, and the NK-cell marker GNLY, were entirely absent from the spectral library, precluding their identification in downstream single cell runs (**Figure 2G**). Conversely, BANK1 and PSAP were present in the sorted B cells and monocytes libraries, respectively, but failed the FDR thresholds during the unified search and were absent from the combined library (**Figure 2G**). Across all sorted cells, canonical markers were detected at widely varying frequencies. The T-cell markers CD3D, CD4, and CD8A were detected in 22%, 7%, and 7% of cells; the B-cell markers CD74, CD79B, and MS4A1 in 43%, 8%, and 13%; the monocyte-associated markers AIF1, CTSS, FSCN1, and MNDA in 23%, 70%, 25%, and 27%; and the NK-cell markers CD7, FCER1G, and GZMB in 27%, 53%, and 22%. These rates quantify feature sparsity across a mixed SCP dataset and underscore the need for careful preprocessing to ensure that clustering is driven by biological differences.

### Benchmarking identifies preprocessing strategies that preserve biological structure

Using the 1,322 QC-passed cells and 1,504 QC-passed proteins, we systematically evaluated how combinations of normalization (e.g., median-centering vs. probabilistic scaling), imputation (e.g., k-nearest neighbors vs. shifted distribution models), and clustering (e.g., graph-based vs. centroid-based methods) influence signal-to-noise and cell-type separation (**Figure 1A**; **Figure 3**). Probabilistic quotient normalization (PQN) (Dieterle et al. 2006) outperformed both median-centering (Vanderaa and Gatto 2023; Leduc et al. 2024) and variance stabilizing normalization (VSN) (Huber et al. 2002), achieving the highest overall quality score, reducing technical and cell volume biases while maximizing cluster quality (**Figure 3A-F**). Eleven distinct clusters were identified through initial exploratory Louvain clustering (**Figure 3G**). Projecting these by cell type and acquisition batch showed that the underlying biological signal was dominant (**Figure 3H**), but batch effects persisted as nested subclusters within each lineage (**Figure 3I**). We therefore applied the Harmony algorithm (Korsunsky et al. 2019) to reduce batch-associated structure while preserving lineage separation (**Figure 3J**). Specifically, distinct cell-type-driven effects were observed in 80% of the cells (clusters 0, 1, 2, 3, 5, 7 and 9), which comprised 74-99% of their respective clusters, while the remaining 20% (clusters 4, 6 and 8) represented mixed populations containing a blend of all four cell types (**Figure 3K**, **Figure S3**).

**Figure 3.**
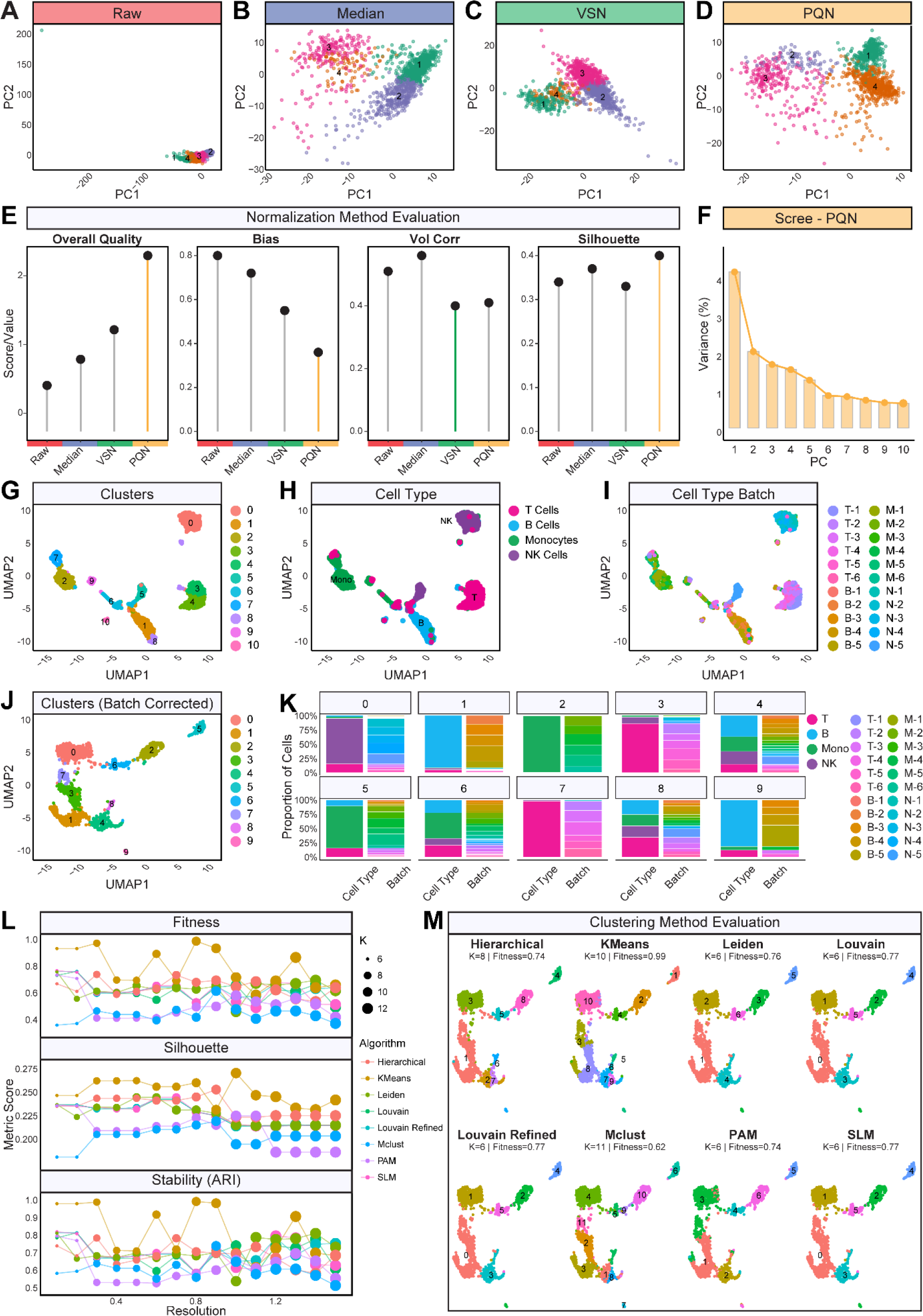
Benchmarking normalization and clustering methods. **A-D**. PCA of sorted cohort under raw, median-centered, VSN, and PQN normalization. **E**. Normalization method evaluation. **F**. Scree plot for the PQN-normalized data. **G**. Initial clustering (Louvain) projected by cluster, cell type (**H**), and acquisition batch (**I**). **J**. Clustering after Harmony batch correction. **K**. Cluster composition per cell type and batch. **L**. Evaluation of eight clustering methods across resolutions. **M**. UMAP visualization of each algorithm at its highest overall quality score.

We next benchmarked eight clustering algorithms: K-means (Sinaga and Yang 2020), hierarchical (Murtagh and Contreras 2012), Leiden (Traag et al. 2019), Louvain (Blondel et al. 2008), Louvain-refined (Rotta and Noack 2011), MClust (Scrucca et al. 2016), partitioning around medoids (PAM) (“Partitioning Around Medoids (Program PAM)” 1990) (Kaufman et al., 1990), and smart local moving (SLM) (Waltman and van Eck 2013) across a multi-metric framework evaluating silhouette width, adjusted rand index (ARI) for cluster stability, and an integrated overall-fitness score. Although K-means (k=10) achieved the highest fitness score, a resolution-dependent sensitivity analysis revealed that its performance was highly unstable, with ARI fluctuating sharply between steps (**Figure 3L-M**). The graph-based clustering methods Leiden and Louvain behaved more predictably, with cluster stability improving as resolution increased above 1.0. Because negative-selection sorting carries along closely related and co-purified subpopulations, we expected the data to contain several distinct clusters (**Figure S2A**). Louvain reached its best fitness score at k=6, but a higher resolution (Res=1.6, k=13) was more stable at a comparable fitness score (**Figure 3L-M**). This rising stability at higher resolutions reflects real biology rather than overclustering, allowing us to isolate co-purified subpopulations and distinct activation states. Centroid-based methods K-means and hierarchical clustering were less stable across resolutions but useful in a different way. They separated out discrete technical anomalies and contaminants that the graph-based methods absorbed into larger clusters. Because graph-based partitioning provided a more stable and biologically coherent framework for continuous immune-cell states, it was selected over the top-scoring K-means model for downstream annotation and classifier training.

### Lineage-resolved SCP identifies immune-cell states in enriched PBMC fractions

To comprehensively resolve intra-lineage heterogeneity, the four major cell types were analyzed independently. We applied stringent per-lineage QC thresholds alongside targeted cell-enrichment strategies (**Figure S2**) to support high-confidence annotation of 1,031 cells (78% of the QC-passed cohort). The remaining 291 cells (22%) were withheld from independent subtyping to protect annotation confidence. Cells that fell below these strict lineage-specific thresholds, or whose proteomic signatures indicated co-purified or cross-lineage identity, were retained and resolved later against the global sorted reference (**Figure 4M**). Unsupervised clustering resolved distinct functional states within each compartment, uncovering four T-cell subpopulations across 304 cells (naïve, memory/migratory, cytotoxic, and proliferating; **Figure 4A**); four B-cell clusters across 268 cells (naïve, proliferating, transitioning, and effector/migratory; **Figure 4D**); three monocyte subsets across 264 cells (proliferating, classical, and non-classical; **Figure 4G**); and three NK-cell states across 195 cells (effector, adaptive, and proliferative; **Figure 4J**).

**Figure 4.**
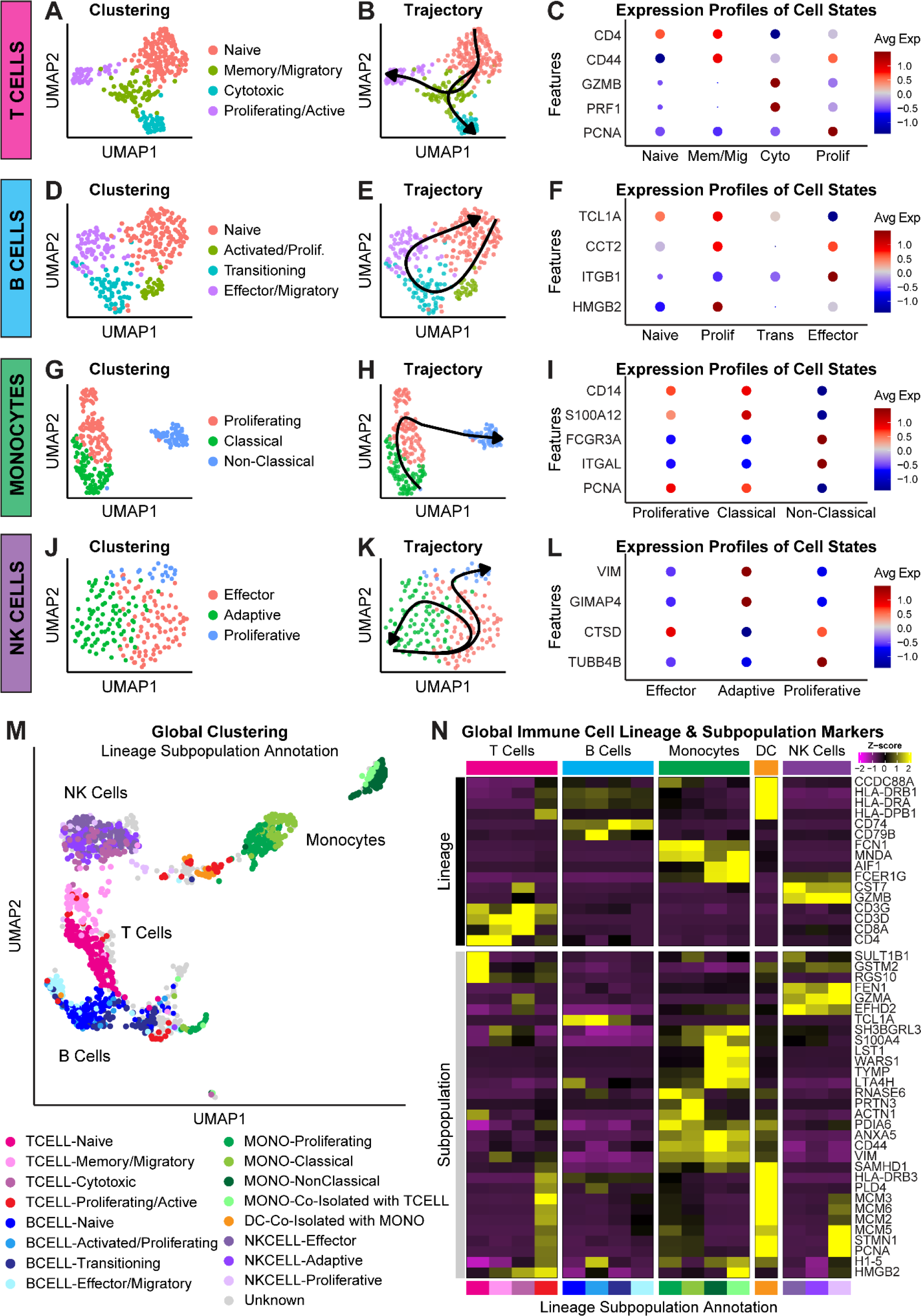
Lineage-resolved subpopulations and integrated global mapping. **A-C**. T cell clustering, trajectory, and marker abundance per state. Same for B cells (**D-F**), monocytes (**G-I**), and NK cells (**J-L**). M. Global UMAP of all sorted cells with lineage subpopulations annotated, including co-isolated DCs and Unknown populations. Unknown cells = passed global QC, not subpopulation-specific QC or cell was removed as a co-isolated cell. **N**. Heatmap of lineage- and subpopulation-defining markers across annotated compartments.

Trajectory inference with Slingshot (Street et al. 2018) revealed consistent proteomic continua within each compartment with pseudotime paths matching known immune-state transitions from naïve states toward specialized effector or proliferative phenotypes (**Figure 4B**, **E, H, K**). In the T-cell lineage, CD4 marked both naïve and memory T cells, with the memory population further distinguished by CD44 up-regulation consistent with a migratory or tissue-homing phenotype (**Figure 4C**). Cytotoxic T cells displayed canonical CD8 abundance with enrichment of the cytolytic effectors GZMB and PRF1. Naïve and effector B cells were defined by the restrictive abundance of TCL1A and CCT2/ITGB1, respectively, while transitioning B cells captured an intermediate state characterized by TCL1A loss prior to induction of mature effector proteins. Within the myeloid lineage, both CD14+ classical and FCGR3A+ non-classical monocytes were clearly resolved (**Figure 4I**), and NK effector and adaptive subpopulations were distinguished by differential abundance of CTSD and VIM/GIMAP4, respectively **(Figure 4L**). Notably, the proliferating clusters across lineages displayed cell-type-specific cell cycle signatures. PCNA expression was restricted to proliferating T and NK cells, whereas elevated HMGB2 abundance defined proliferative monocytes and T subsets, and H1-5 accumulation uniquely characterized the dividing B cell cluster (**Figure 4C**, **F**, **I**, **L**, **N**).

### Integrated mapping identifies co-purified populations and annotation limits

We projected the lineage-specific cell identities back onto the global dataset in which all cell types were searched together (**Figure 4M**). These labels revealed clear phenotypic compartmentalization across localized subpopulations, particularly within the monocyte, T, and B cell compartments. Specifically, classical monocytes clustered apart from non-classical monocytes; naïve and memory/migratory T cells split by subpopulation; and naïve B cells clustered independently but adjacent to effector B cells. This global mapping also allowed us to preserve maximum cellular data by rescuing co-purified cell populations that had been flagged during independent QC. Rather than discarding cells displaying cross-lineage markers, populations such as dendritic cells (DCs) co-isolated within the monocyte pool (n=32), or monocyte contaminants within the T cell lineage (n=17), were successfully resolved and mapped onto the global sorted dataset (**Figure 4M**). This approach also enabled refinement of proliferating T cells that had clustered with DCs due to shared protein profiles (**Figure 4N**). Phenotypic reassessment of this subpopulation revealed a distinct cluster with a larger physical diameter, down-regulation of the T-cell receptor signaling protein ZAP70, and high abundance of the myeloid marker SYK. This signature explained the anomalous clustering and allowed accurate re-annotation as DCs co-isolated with T cells (**Figure S4**). Finally, cells rejected by the strict independent lineage filters but retained by the more relaxed global thresholds and not relabeled as a co-isolated cell type were designated as ‘Unknown’ (**Figure S5A**). For example, 142 NK cells with markedly lower protein intensity than the rest of the compartment were excluded from the independent NK analysis to preserve the fidelity of downstream subtyping (**Figure S5B**). Because these cells met the global processing parameters, they were preserved and labeled ‘Unknown’ in the final global layout (**Figure 4M**). Together, the distinct lineage identities and subpopulation signatures (**Figure 4N**) validate the processing pipeline and establish an annotated single-cell immune landscape suitable for training a downstream predictive classifier for automated subpopulation annotation.

### Mixed PBMC analysis validates SCP-based immune-cell classification

Simultaneous analysis of the unsorted PBMC fraction yielded 334 high-quality single cells (**Figure 5A**). Unsupervised clustering identified eight subpopulations without prior algorithmic guidance, each significantly enriched for cell-type-specific proteomic signatures (**Figure 5B-C**). Cluster 0 (20.4% of the dataset) showed robust enrichment for a shared NK/CD8+ T-cell signature driven by cytotoxic markers including CST7. Clusters 1 (19.2%) and 5 (6.9%) were enriched in T- and B-cell markers, respectively, bringing the putative lymphocyte fraction to 46.5%. This lymphocyte fraction falls below 70-90% typically observed in healthy PBMCs. Notably, scRNA-seq performed on individual cells from the same PBMC sample, processed on the same day, recovered quantitatively similar proportions, indicating that this distribution reflects the composition and handling of this preparation rather than an SCP-specific artifact. Cells with an enriched monocyte profile clustered separately from the lymphoid lineages. The spatial organization also reflected developmental biology: monocytes and DCs share foundational lineage markers, producing overlapping signatures within clusters 3 and 4 (**Figure 5C**), while B cells and DCs, both antigen presenting cells (APCs) sharing HLA class II proteins, remained globally distinct, with monocytes and DCs clustering close together and B cells and DCs farther apart in latent space. This lineage-specific segregation validates the pipeline’s capacity to preserve biological signal over technical noise.

**Figure 5.**
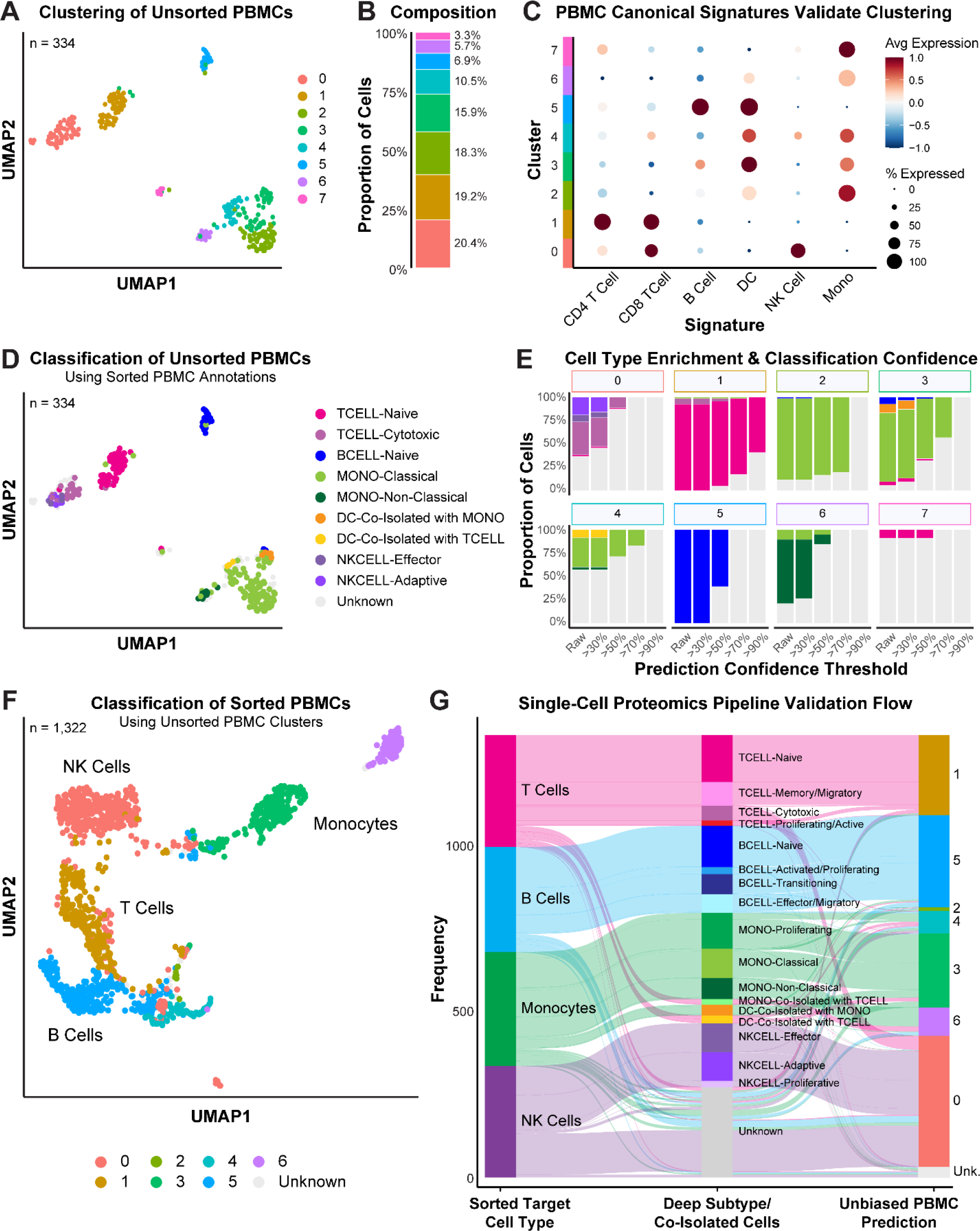
SCP-based classification of unsorted PBMCs and bidirectional validation. **A.** Unsupervised clustering of unsorted PBMCs. **B**. Cluster composition. **C**. Canonical signature scores across clusters. **D**. Unsorted PBMCs classified using sorted-cell annotations. Confidence threshold ≥ 0.5. **E**. Cell type enrichment for each cluster across increasing confidence thresholds. **F**. Sorted cells classified using unsorted PBMC clusters. G. Alluvial plot showing flow from sorted lineage to deep subtype and co-isolated cell identity, to unbiased unsorted PBMC prediction.

To independently validate the unsupervised annotations and achieve higher-resolution phenotypic mapping, we classified the mixed PBMCs using our enrichment-derived lineage labels with a probability threshold of > 50% to ensure high-confidence assignments (**Figure 5D**). The supervised predictions agreed strongly with the unsupervised clustering and parsed broad lineage assignments into functional compartments: naive and cytotoxic T cells, naive B cells, classical and non-classical monocytes, DCs, and effector and adaptive NK cells. To quantify accuracy of the predictions, we evaluated the cell-type enrichment within each cluster across increasing confidence thresholds (**Figure 5E**). Naive CD4+ T cells had the most distinct proteomic signature, demonstrating high separability and robust confidence prediction. Over 50% of the naive T cells in cluster 1 were predicted with ≥ 90% confidence. Similarly, classical monocytes were predicted with high confidence, with over 75% cluster-2 cells exceeding 70% confidence. DCs, restricted to clusters 3 and 4, showed lower confidence, likely because of their small sample size (n = 43). These clusters also contained a substantial fraction of classical monocytes. Highly specific enrichment was observed for non-classical monocytes, which isolated cleanly into cluster 6, and B cells to cluster 5. Even with only 334 cells, we resolved distinct functional subtypes rather than just major lineages. Scaling this approach to larger numbers should allow for even higher-resolution subtyping in PBMC samples.

This biological distinctness was further substantiated by an independent cross-validation analysis across seven classification methods: support vector machine (SVM) (Cortes and Vapnik 1995; Karatzoglou et al. 2004), weighted random forest (RF) (Breiman 2001; Ishwaran et al. 2008), balanced random forest (BRF) (C. Chen 2004; Wright and Ziegler 2017), gradient-boosted trees (LightGBM) (Ke et al. 2017; Friedman 2001), k-nearest neighbor (KNN) (Cover and Hart 1967), and a single- and multi-layer perceptron neural network (NN_SLP, MLP) (Rumelhart et al. 1986) (**Figure S6**). Non-classical monocytes and naive T cells ranked as the two most easily classified subpopulations, achieving the highest cumulative F1-scores, confirming their unique proteomic signatures. The lowest-ranked were the proliferating subsets (NK-, T-, and B-proliferative; all n <30), consistent with the mixed-PBMC classification in which no cell was predicted as proliferating (**Figure 5D**). Although cell-type-specific cell cycle markers were observed for the proliferating cells, they were insufficient for robust classification likely because of shared proliferation and lineage signatures among the training labels. Notably, BRF surfaces the proliferating cells but at a precision cost (**Figure S6**).

### A bidirectional transfer learning approach accurately decodes the cellular heterogeneity of unsorted PBMCs

To test whether our feature selection operates without algorithmic bias, we performed the reverse validation. We extracted the independently generated, unsupervised cluster labels from the unsorted PBMCs and used them to classify the 1,322 cells of the global sorted reference (**Figure 5F**). The concordance between forward (sorted → mixed) and reverse (mixed → sorted) classification provides strong evidence that the recovered lineage structure reflects genuine proteomic identity rather than an artifact of the clustering applied to either dataset. This bidirectional correspondence is summarized as a flow from sorted target lineage, through deep subtype and co-isolated identity, to the unbiased unsorted-PBMC cluster prediction (**Figure 5G**). The sorted lineages branch into finer subtypes that map cleanly onto our unsorted PBMC clusters, keeping the lineage identities intact across the pipeline. Together, these results demonstrate that SCP can resolve complex PBMC structures strictly on its own, without relying on scRNA-seq guidance or proxy-labeling.

## DISCUSSION

We combined cell sorting, cellenONE dispensing, and Orbitrap Astral Zoom acquisition at 70 SPD with a data-driven analytical pipeline to profile human PBMCs by SCP. From 2,427 single-cell proteomes, we established a QC-filtered reference of 1,322 sorted cells, benchmarked preprocessing and clustering choices, resolved functional immune-cell states within each lineage, and validated the resulting annotations bidirectionally against an unsorted PBMC mixture. Beyond the annotated landscape itself, the pipeline provides analytical choices, including ambient background modeling, normalization, and clustering method selection, that can be applied to other heterogeneous/immune SCP studies. Our approach builds on the growing ecosystem of SCP analysis tools such as QuantQC (Leduc et al. 2024) and the scp Bioconductor infrastructure (Vanderaa and Gatto 2025).

The negative-selection sorting, which was chosen to preserve native proteomes by avoiding surface receptor engagement (Hornschuh et al. 2022), inevitably co-purifies related and interacting cell types. Rather than treating these as contamination to be discarded, our global mapping strategy rescued and correctly re-annotated these populations (i.e., DCs traveling with monocytes and T cells, and monocyte contaminants within the T-cell fraction) based on their distinct proteomic signatures, only identifiable at single-cell resolution. A cluster initially labeled proliferating T cells, for example, was reclassified as co-isolated DCs from its enlarged diameter, loss of ZAP70, and gain of SYK. Because SCP measures the effector molecules of cell identity directly, it can resolve co-purified and transitional populations that share lineage markers but differ in their global proteomes. Simultaneously, our data shows that single canonical markers are unreliable in current MS-based SCP. Many lineage-defining proteins were absent from the spectral library, were removed due to quality issues, or fell below detection. Classification and reannotation were therefore most robust when driven by global proteome patterns rather than individual markers.

Lastly, our benchmarked, classifier-ready SCP workflow for PBMCs provides a natural template for profiling patient samples, where protein-level states such as activation, cytotoxicity and exhaustion are directly correlated to the pathology and treatment response. Applying this pipeline to diseased cohorts would test whether the immune-cell proteomic states defined in healthy PBMCs are remodeled in disease.

## Acknowledgements

We acknowledge the support of the IMS at City of Hope Comprehensive Cancer Center supported by the National Cancer Institute of the National Institutes of Health under award number P30CA33572. We thank Joshua Cantlon of Scienion for discussions and constructive comments regarding the cellenONE and single-cell proteomics sample preparation. We thank the TGen Collaborative Sequencing Center in Phoenix, AZ for sequencing our RNA library on their Illumina NovaSeq X platform. This work was funded by a donation from the Lennar Foundation

## METHODS

### PBMC Samples

Cryopreserved human PBMCs were purchased from Sanguine Biosciences from a 47-year-old healthy male donor (Participant ID: 139193) with no history of infectious disease, tobacco, alcohol, or recreational drug use. Donor PBMCs were collected at a single time point and aliquoted for use in individual experiments by the commercial vendor.

### Sample Thawing and Viability Assessment

Frozen PBMCs were thawed and cell viability and concentration were assessed as previously described (Disis et al. 2006). All centrifugation steps during thawing were performed at 300g for 10 min. Cells were maintained on ice or at 4°C unless otherwise stated.

### Magnetic-Activated Cell Sorting

Miltenyi Biotec cell negative-selection isolation kits were used to sort T cells, B cells, NK cells, and monocytes following the vendor protocols for manual magnetic labeling. After isolation, sorted cells were collected, centrifuged at room temperature, and resuspended in DPBS. For consistency, unsorted PBMCs were processed analogously to sort T, B, and NK cells excluding the use of antibody cocktails and microbeads. Cell viability and concentration of resuspended sorted and unsorted cell pellets were assessed as outlined above. A fraction of unsorted PBMCs were used for scRNA-seq. The remaining sorted and unsorted PBMCs were used for proteomics.

### Sample preparation for single-cell proteomics

For dead-cell labeling, 3 µL/mL SYTOX Green Nucleic Acid Stain (Thermo Scientific, S7020) was added to resuspended sorted and unsorted cells in DPBS for 20 min. Cells were centrifuged to remove residual stain, and cell pellets were resuspended in 1× PBS (PBS; Fisher Scientific, 10-010-031). For single-cell isolation on the cellenONE X1, cells were diluted with 1× PBS to a concentration of 300 cells/mL. To monitor background contamination, negative-control wells (10–12 per 384-well plate) were processed identically excluding cell isolation.

### Sample preparation for single-cell RNA sequencing

Unsorted PBMCs were centrifuged to remove DPBS then cell pellets were washed then resuspended with PBS containing 0.04% ultrapure bovine serum albumin (Thermo Scientific, AM2616). scRNA-seq libraries were prepared using the 10x Genomics Chromium GEM-X 3’ v4 kit (10x Genomics, GEM-X Single Cell 3’ v4 CG000731 Rev B) according to the manufacturer’s protocol. cDNA concentration and quality were determined after amplification using the High Sensitivity D5000 ScreenTape on an Agilent 4200 TapeStation. 25% of total cDNA was used as input for RNA library preparation. The library was diluted at 1:25 in Tris-EDTA buffer and sequenced on an Illumina NovaSeq X platform targeting a depth of 25,000 reads per cell. Raw FASTQ files were generated for downstream single-cell analysis. Raw sequencing data pre-processing and reference genome alignment were done using Cell Ranger.

### Single-cell proteomics cell isolation, digestion, and peptide drying

T cells, B cells, NK cells, monocytes, and unsorted PBMCs were dispensed via the cellenONE X1 platform into 384-well plates preloaded with lysis buffer (100mM triethylammonium bicarbonate buffer-0.2% n-dodecyl-beta-D-maltoside) which were then frozen at -80°C and lysed as previously described (Petelski et al. 2021). After denaturation, 10 ng of degassed mass spectrometry-grade Trypsin Gold (Promega, V5280) was dispensed into 384-well sample plate samples which were then incubated at 37 °C, acidified with 0.1% formic acid, dried via a SpeedVac centrifuge (Eppendorf, 022820168), and then frozen at -80°C for storage until LC-MS/MS acquisition.

### Liquid chromatography–mass spectrometry

Frozen 384-well sample plates containing SCP samples were thawed, and SCP sample peptides were then reconstituted in 0.015% n-dodecyl-beta-D-maltoside-0.1% formic acid containing Pierce Peptide Retention Time Calibration Mixture (Thermo Scientific, 88320). Peptides were separated on an 8 cm C18 column (IonOpticks Aurora AUR4-25075C18-XT, 75 µm ID, 1.7 µm particle, 120 Å pore size) heated to 50°C. Samples were directly injected onto the column at a pressure of 1350 bar via a Vanquish Neo UHPLC System (Thermo Fisher Scientific) and nanosprayed using a spray voltage of 1.8 kV into Orbitrap Astral Zoom mass spectrometer equipped with a FAIMS Pro device (FAIMS CV -48) and the ion transfer tube maintained at 280 °C. The carrier gas flow was set to 3.5 L/min. Peptides were eluted using a variable flow 11 min active gradient formed by Solvent A (LC-MS grade water, 0.1% formic acid) and Solvent B (80% LC-MS grade Acetonitrile, 20% LC-MS grade water, 0.1% formic acid). Briefly, the gradient composition was 4%, 12%, 22.5%, and 40% B from 0.0-0.1, 0.1-1.7, 1.7-6.7, and 6.7-8.7 min, respectively, with the remaining run time consisting of 99% B to wash the UHPLC column.

Data-independent acquisitions were performed on a Thermo Scientific Astral Zoom mass spectrometer using the predefined low-input single-cell template in Xcalibur Method Editor (Thermo Fisher Scientific Xcalibur software version 4.7.102.25). Full MS1 scans were acquired with the Orbitrap mass analyzer, and MS2 scans were acquired with the Astral mass analyzer. The data were acquired with a cycle time of 80 ms. Thermo Fisher Scientific Tune software version 2.1.765.17 was used to acquire MS data.

### Data Analysis

Raw data were searched using Spectronaut (version 20.5.260227.92449, Biognosys) via directDIA mode (Bruderer et al. 2015). Data files were searched in MS run batches without cross-normalization with every raw file defined as a separate condition. Libraries were created using MS run batches, and quantification was performed at the MS1 level. Cysteine carbamidomethylation was removed as a static modification. Default settings were used for all analyses and for library generation unless otherwise indicated. Searches were performed against the human proteome (UniProt proteome UP000005640, reviewed, 42,518 entries, downloaded December 2025). FDR filtering was based on Spectronaut default settings and 1% at the protein level. For FDR checks, a decoy (‘shuffled target’) database was generated with fixed positions for arginine, proline and lysine.

### Quality Control and Downstream Analysis

Single-cell proteomics data were processed through a custom in-house pipeline implemented in R. cellenONE isolation metadata were mapped to their corresponding single-cell proteomics measurements, and cells were assessed against quality-control metrics including protein identification, intensity, and data completeness. Cells passing QC were retained for downstream normalization, dimensionality reduction, and clustering. Batch-level comparisons were evaluated using non-parametric statistical tests. All analyses were performed in R (version 4.4.2).

